# Stressed Avoider rats show blunted sensitivity to alcohol’s aversive effects: Potential contributions of the lateral habenula and lateral hypothalamus

**DOI:** 10.1101/2025.05.23.655819

**Authors:** OR Brunke, SM Bonauto, EM Boshak, MJ Rauch, MM Weera

**Affiliations:** Department of Psychology, Tufts University, Medford, MA; Neuroscience Program, Graduate School of Biomedical Sciences, Tufts University, Boston, MA

**Keywords:** avoidance, aversion, predator odor, alcohol, lateral hypothalamus, lateral habenula, central amygdala

## Abstract

Avoidance coping following stress exposure predicts heightened alcohol drinking. Similarly, blunted sensitivity to the aversive effects of alcohol facilitates increased drinking. However, the relationship between stress exposure, coping mechanism, and sensitivity to alcohol’s aversive effects is unknown. In rats, predator odor stress increases alcohol intake in animals that show persistent avoidance of stress-paired stimuli, termed “Avoiders”. Here, we tested the hypothesis that Avoider rats have blunted sensitivity to alcohol’s aversive effects using an alcohol-induced conditioned taste aversion (CTA) paradigm. After a single conditioning session, Non-Avoider rats acquired alcohol-induced CTA while Avoiders did not. Male rats across all groups eventually acquired alcohol CTA after four conditioning sessions. However, in females, only Non-Avoiders acquired alcohol-induced CTA. In male Non-Avoider rats, a single CTA-inducing dose of alcohol increased cFos expression in the lateral habenula (LHb), an important nucleus in aversion signaling. In male Avoiders, the same dose of alcohol decreased LHb cFos expression. cFos expression in the lateral hypothalamus (LH), which provides glutamatergic inputs to the LHb, was also diminished by alcohol in male Avoider rats. In females, alcohol had no effect on cFos cell counts in the LHb. However, in the LH, alcohol diminished cFos expression in female Non-Avoiders. Collectively, these findings suggest that stressed Avoider rats are hyposensitive to alcohol’s aversive effects, which may facilitate their heightened alcohol drinking after stress. Sex- and stress group-specific differences in LH and LHb recruitment highlight these regions as candidates for mediating stress-induced changes in alcohol behaviors.

**Highlights:**

- Avoider rats exhibit blunted sensitivity to alcohol’s aversive effects
- Alcohol dose increases cFos expression in the lateral habenula of male Non-Avoiders
- Alcohol dose diminishes cFos expression in the LHb and LH of male Avoiders

## 1. Introduction

Exposure to stressful events may lead to adverse consequences such as increased anxiety, depression, as well as alcohol and substance misuse. Excessive alcohol drinking bears a substantial socioeconomic toll, contributing directly to ∼178,000 deaths (CDC) and inflicting an economic cost of ∼$250 billion per year in the United States alone (Sacks et al., 2015). Studies show that individuals who respond to stress with avoidance coping (i.e., evasion rather than active confrontation of stressors and stress-associated stimuli) are more likely to escalate their alcohol drinking after stress (McKee et al. 1998; Hasking & Oei 2007; Bartone et al. 2017). Similarly, in people diagnosed with post-traumatic stress disorder (PTSD), avoidance symptoms predict not only more severe PTSD outcomes, but also higher levels of alcohol misuse (Hruska et al. 2023). Collectively, empirical evidence in humans indicates an important relationship between avoidance stress coping and alcohol misuse.

In lab rats, exposure to predator odor stress (i.e., bobcat urine) produces avoidance behavior in only a subset of animals, termed ‘Avoiders’, mirroring individual differences in stress responsivity in humans. In this model, all predator odor-exposed animals show evidence of acute stress, such as elevated serum adrenocorticotropic hormone (ACTH) and corticosterone in the hours after predator odor exposure, blunted body weight gain, and heightened anxiety-like behavior over a period of several days after stress (Whitaker & Gilpin, 2015; Weera et al., 2020). However, only Avoider rats show conditioned avoidance of predator odor-associated contexts that remains stable over weeks after stress and is not altered by repeated odor exposure (Schreiber et al., 2017; Weera et al., 2020). Predator odor stress also heightens alcohol consumption in Avoider, but not in Non-Avoider rats. Specifically, Avoider rats show escalated operant alcohol self-administration and free-choice drinking after predator odor exposure compared to Non-Avoiders and unstressed Controls, as well as to their own pre-stress baseline (Edwards et al., 2013; Weera et al., 2021; Bonauto et al., 2025). These self-administration studies were initially performed in male rats but were recently replicated in both males and females (Weera et al., 2023).

Alcohol drinking, particularly in recreational or social contexts, is typically viewed as being driven by alcohol’s hedonic, positive reinforcing properties. On the other hand, alcohol drinking by alcohol-dependent individuals is primarily motivated by negative reinforcement (Koob 2013a, 2013b, 2014). In addition to its positive and negative reinforcing properties, alcohol has inherent aversive properties and activates brain circuits that signal aversion (Glover et al., 2016; Pryzybysz et al., 2024; Cunningham et al., 2003, 2019, 2021; Riley et al., 2022). Studies in humans show that a lower subjective sensitivity toward alcohol’s aversive effects predicts higher levels of alcohol drinking (Schuckit 1994, 2014), indicating that alcohol’s aversive effects, particularly at higher doses achieved through drinking, serve as a potent regulator of alcohol intake. Indeed, laboratory studies in which human subjects were challenged with various doses of alcohol show that individuals who were more sensitive to alcohol’s aversive effects (e.g., subjective ratings, sedation) were less likely to binge drink and to develop alcohol use disorder (AUD) at future time points (King et al. 2011, 2019). Similarly, in rodent models, studies show that animals that are less sensitive to the aversive properties of alcohol, as measured using assays such as alcohol conditioned taste aversion (CTA) or place aversion (CPA), tend to drink or self-administer more alcohol (Chester et al. 2003; Cunningham et al. 2019). Therefore, rodent models are useful for elucidating mechanisms of alcohol aversion, which may eventually be leveraged for the management of alcohol misuse and AUD.

In the brain, the habenula is an epithalamic nucleus that plays a key role in aversion signaling (Matsumoto & Hikosaka 2007, 2009; Hikosaka 2010). The habenula receives sensory and visceral information from the hypothalamus, thalamus, and various forebrain regions, and sends projections to midbrain areas such as the ventral tegmental area (VTA) and rostromedial tegmentum (RMTg) (Jhou et al. 2009; Balcita-Pedicino et al., 2011; Lecca et al. 2017). Lateral habenula (LHb) neurons are alcohol responsive, such that even low doses of ethanol increase action potential firing in these cells (Zuo et al. 2017). With regards to behavioral control, LHb activation in response to alcohol signals aversion, as measured via alcohol CTA (Haack et al. 2014; Glover et al. 2016) and CPA (Zuo et al. 2017). In line with the idea that alcohol’s aversive properties are important for limiting consumption, lesions of the LHb are sufficient to increase alcohol drinking, operant self-administration, and physiological stress-induced reinstatement of alcohol seeking (Haack et al. 2014).

Given that Avoider rats increase their alcohol consumption after stress, the goal of this study was to test the hypothesis that stress blunts alcohol aversion and alcohol-induced LHb activation in Avoider rats. Using an alcohol taste conditioning procedure, we found that a single alcohol challenge was sufficient to produce alcohol CTA in Non-Avoiders, but not in Avoider rats. Using immunohistochemistry, we found that alcohol challenge increased the number of cFos-positive cells in the LHb of Non-Avoiders but decreased the number of cFos cells in the LHb of Avoider rats. Alcohol also decreased cFos-positive cells in the lateral hypothalamus (LH), a brain area that sends dense glutamatergic projections to the LHb, in Avoiders.

## 2. Materials & Methods

### 2.1 Animals

Male and female Wistar rats used in the following experiments arrived at 8 weeks of age (Charles River, Kingston, NY) and were pair housed with same-sex cage mates in a temperature-and humidity-controlled vivarium on a 12-hour light/dark cycle (lights off at 0900). Food (Teklad 2919) and tap water were available *ad libitum* except where described. All behavioral tests occurred in the dark period. All procedures were approved by the Tufts University Institutional Animal Care and Use Committee (IACUC).

### 2.2 Predator odor place conditioning

#### Place Conditioning Procedure

Rats were subjected to a 4-day predator-odor place conditioning paradigm (Edwards et al., 2013, Weera et al., 2023, Bonauto et al., 2025). On day 1, rats were allowed to explore a three-chamber apparatus with a middle zone and two distinct contexts (mesh floors with striped walls vs holed floors with dotted walls; Pretest). The duration in each context was quantified. On day 2, individual rats were confined to either context for 15 minutes without the presence of any unconditioned stimuli (Neutral Conditioning). On day 3, rats were confined to the opposite chamber in the presence of a bobcat urine-soaked paper towel placed on a tray underneath the mesh or holed floors for 15 minutes (Odor Conditioning). Unstressed Controls were exposed to similar handling but never to predator odor. At the end of the Odor Conditioning day, the apparatus was cleaned with soap and water and deodorizing spray was liberally utilized on all surfaces (PureAyre Odor Eliminator, Clean Earth Inc., Kent, WA). On day 4, individual rats were again allowed to explore the entire arena and duration in odor-paired chamber was quantified (Posttest). Odor-exposed rats that avoided the odor-paired context for greater than 10 seconds compared to their individual baseline during the Pretest were considered Avoiders and other odor-exposed rats were classified as Non-Avoiders.

#### Additional behavioral analysis

Videos were analyzed in Ethovision XT (v17.5, Noldus Information Technology) for a variety of behaviors on the Posttest day of predator odor CPA. Fifteen videos were excluded due to poor video quality that inhibited automated behavioral tracking (excluded videos were balanced across groups). Latency to enter the odor-paired context, number of entries into the odor-paired context, and percentage of time in the center zone (proximal to odor context) were quantified and compared between groups of stressed animals (Avoiders vs Non-Avoiders). Behaviors were analyzed separately in males and females due to established differences in avoidance and anxiety-related behaviors (Gruene et al., 2015).

### 2.3 Alcohol-induced conditioned taste aversion

Rats (60 male and 60 female) were left undisturbed for 4 days following a Thursday arrival and were subsequently handled on 4 consecutive days (Monday-Thursday) prior to habituation saline injections (10 ml/kg, i.p.) on Friday (Figure 1A). Beginning the following Monday, rats were water-restricted for 6 days, such that they only had water access for 30 min per day beginning 30 min into the dark cycle (0930) in a drinking chamber distinct from their home environment. Rats were pair-housed throughout the experiment except during this 30 min period of fluid consumption. On Tuesday-Friday of this week, rats also underwent the predator odor CPA procedure and indexed for avoidance as described above (starting 3-4 hours into the dark cycle). Beginning on a Monday of the following week, instead of water, rats were given a novel 0.1% saccharin solution for 30 min in their respective drinking chambers. By this time, rats were conditioned to consume large volumes of fluid during their daily hydration window. At the end of this 30 min saccharin intake window, rats were injected (i.p.) with either 1.5 g/kg alcohol (in 0.9% saline) or an equivalent volume of saline (10 ml/kg). This procedure was repeated 48h later (i.e., on Wednesday), and the change in intake of the novel saccharin-flavored solution from Monday (Session 1) to Wednesday (Session 2) was used as a measure of the conditioned effects of acute alcohol. This procedure was repeated for two additional sessions separated by 48h (i.e., on Friday and Sunday) to generate alcohol taste conditioning acquisition curves. On the intervening days (Tuesday, Thursday, and Saturday), rats were given plain water to maintain hydration. Rats were weighed daily to ensure adequate health. Rats that consumed less than 1ml of saccharine during the first session were excluded from analysis due to lack of exposure to the conditioned stimulus.

**Figure 1:**
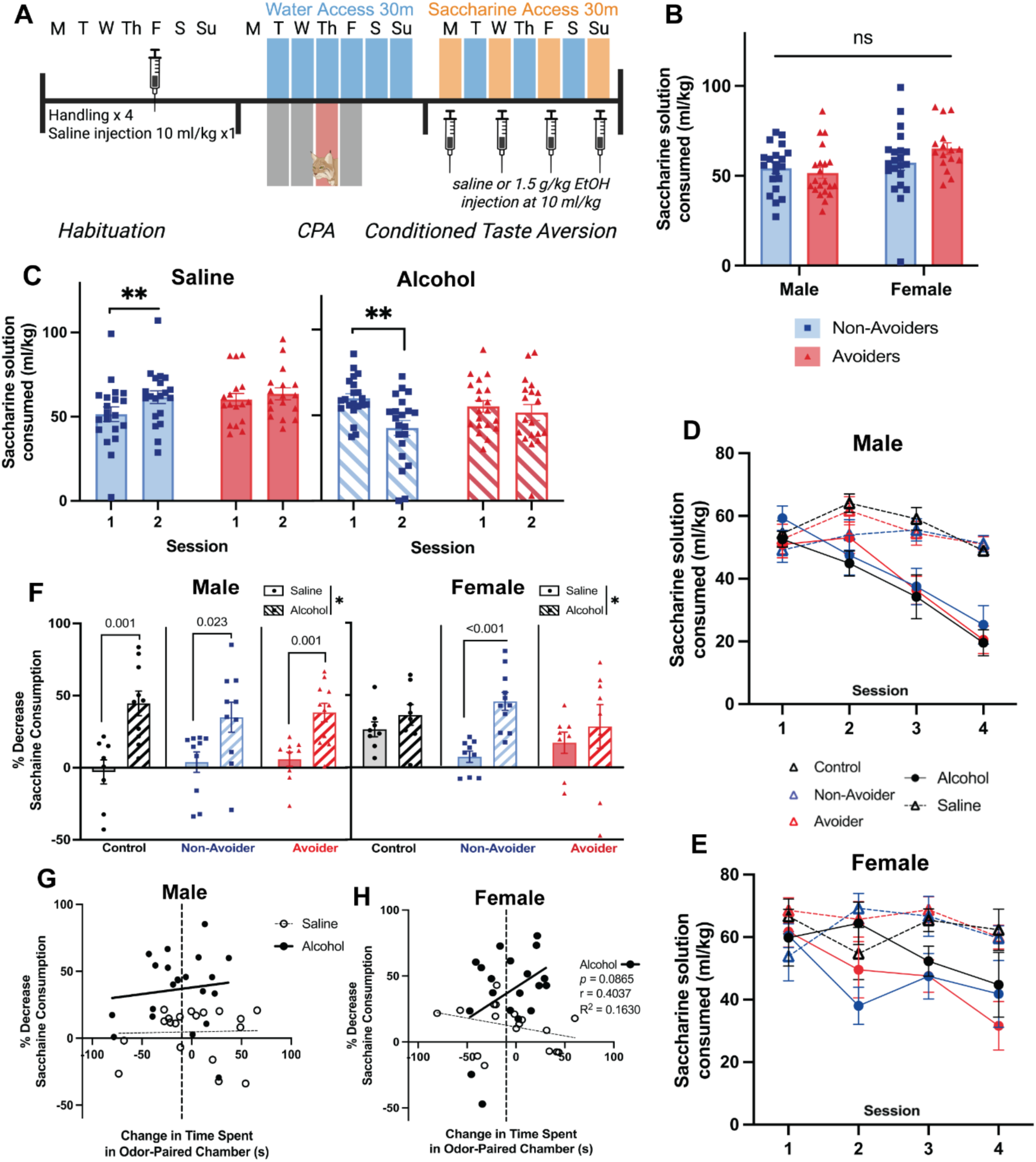
Non-Avoider rats develop CTA to 1.5 g/kg alcohol injection, some Avoiders do not. **A.** Experiment timeline. **B.** Pre-conditioning baseline saccharin intake (in Session 1) in Avoiders and Non-Avoiders. **C.** Change in saccharin intake after one conditioning session in which an alcohol or saline injection was paired with saccharin access. **D.** Saccharin intake in each of all 4 conditioning sessions in male and **E.** female rats. **F.** Percent decrease in saccharin consumption from the first to the last conditioning session. **G.** Correlations between time spent in odor-paired chamber (avoidance scores) and percent decrease in saccharin consumption in males and **H.** females. Created in BioRender. Bonauto, S. (2025) https://BioRender.com/klnmq1f

### 2.4 Blood alcohol concentrations following alcohol injection

A separate cohort of 12 (6 male and 6 female) Wistar rats were housed under similar conditions. Blood samples were collected via tail snips from each rat (11 weeks old) immediately before alcohol injection (i.p.; 20% ethanol in 0.9% saline), and at 30 minutes and 90 minutes post-injection. Rats were given alcohol doses of either 0.5 g/kg, 1.5 g/kg, or 2.0 g/kg. Samples were centrifuged to isolate serum no more than 1-hour post-collection and serum was stored at -80°C until blood alcohol concentration (BAC) measurement using an AM1 Alcohol Analyer (Analox, Stourbridge, UK).

### 2.5 cFos expression following alcohol challenge

#### Alcohol challenge and tissue collection

Rats (32 females, 35 males) were handled and subjected to predator odor place conditioning as described above. One day after the predator odor place conditioning procedures were completed (i.e., 48h after stress exposure), rats were injected with 1.5 g/kg alcohol or an equivalent volume of saline in the home cage. Animals were euthanized 90 minutes later with 5% isoflurane and brains were post-fixed. Half of the animals were transcardially perfused with phosphate-buffered saline and 4% paraformaldehyde (PFA), and their brains were post-fixed in 4% PFA overnight. The other half of the rats were euthanized without perfusion, and their brains were post-fixed in Zinc Formalin fixative (Sigma Aldrich, catalog #Z2902) for 48 hours. All brains were cryoprotected in 20% sucrose, snap frozen on dry ice, and stored at -80 °C until cryostat sectioning.

#### Immunohistochemistry

Brains were sectioned coronally at 40 µm using a cryostat, and the sections were stored free-floating in 1X phosphate buffered saline (PBS) containing 0.1% sodium azide at 4°C. Every sixth section from bregma -1.5 to -3.9 was processed for immunohistochemistry (IHC) to stain for c-Fos. All steps were completed at room temperature unless otherwise stated. First, sections were incubated in 1% hydrogen peroxide in 1X PBS to quench endogenous peroxidase activity. Then they were blocked via incubation in 5% normal donkey serum for 90 minutes. At the end of day one, sections were incubated in the primary antibody to label c-Fos (1:1000; Cell Signaling Technology, Danvers, MA) in 1X PBS, 0.5% TWEEN-20, and 5% normal donkey serum) overnight at 4°C. Next sections were washed with 1X PBS and incubated in horse-anti-rabbit horseradish peroxidase Immpress Kit (Vector Laboratories, Burlingame, CA). Finally, sections were incubated in a DAB Substrate with nickel kit (Vector Laboratories, Burlingame, CA) to develop the signal. Sections were then mounted onto slides, dehydrated via ethanol bath, and coverslipped with Permount.

#### Imaging and quantification

Images were captured using the Keyence BZ-X800 Microscope (Keyence Corporation, Osaka, Japan) and cFos positive cells were counted within each region using Fiji software (Schindelin et al., 2012). The lateral hypothalamus (bregma -1.56 to -2.64), lateral habenula (bregma -2.92 to -3.84), and central amygdala (bregma -1.56 to -2.92) were imaged at 20x magnification and the number of cFos positive cells per unit area (pixels) was expressed as cFos density for each brain region. Cell counts were performed by a researcher blinded to experimental conditions.

### 2.6 Statistical analysis

Statistical analysis was performed using SPSS Statistics Version 29 (IBM) and graphs were made using Prism 10 (GraphPad). Data was analyzed using omnibus ANOVAs with Sex, Stress Group, and Treatment as between-subjects factors. When significant interactions were found, lower-order ANOVAs and Tukey’s post-hoc tests were conducted. For CTA, repeated measures ANOVAs were used to test within-subject changes for each sex (Session x Stress Group x Treatment). Pearson correlations were performed and analyzed with simple linear regression analyses. Outliers in t-distribution of each behavioral measurement were detected using the Grubbs Outlier Test and removed only from the corresponding analysis. All data are presented as Mean ± SEM.

## 3. Results

### 3.1 Avoider rats show blunted sensitivity to an aversive dose of alcohol

The goal of this study was to test Avoider and Non-Avoider rats’ initial subjective sensitivity to alcohol’s aversive effects using alcohol taste conditioning. Rats were exposed to predator odor stress, indexed for avoidance, and subjected to an alcohol taste conditioning procedure beginning 4 days post-stress (Figure 1A). Specifically, Avoider and Non-Avoider rats were allowed to consume a novel tastant (0.1% saccharin) for 30 minutes and were challenged with 1.5 g/kg alcohol or saline at the end of this drinking session. This procedure was repeated every 48 hours for a total of 4 saccharin (CS)-alcohol taste conditioning sessions. Importantly, Avoider and Non-Avoider rats showed similar levels of saccharin intake in Session 1 (i.e., pre-conditioning baseline before alcohol challenge; *p* = 0.844) (Figure 1B).

After a single alcohol challenge, Non-Avoider rats significantly reduced their intake of the novel tastant from Conditioning Session 1 to 2, indicating development of alcohol CTA (Repeated Measures Session x Stress Group x Treatment interaction, *F_1,68_* = 3.575, *p* = 0.063; effect of Session in Non-Avoiders, *F_1,20_* = 11.911, *p* = 0.003; Figure 1C). However, this effect was absent in Avoider rats (*p* = 0.344), indicating that Avoiders are less sensitive than Non-Avoiders to the initial aversive effects of alcohol. Interestingly, Non-Avoider rats that were given a saline injection at the end of Session 1 increased their saccharin intake from Sessions 1 to 2 (*F_1,18_* = 17.416, *p* < 0.001), potentially as a result of habituation to the novel tastant, whereas the slight increase in saccharin intake in Avoiders was not significant (*p* = 0.250). There were no significant main or interactive effects with sex, therefore males and females were combined for the above analyses.

With additional CS-alcohol pairings across 4 conditioning sessions, male and female rats developed distinct patterns of conditioned taste aversion (Repeated Measures within-subject Session x Sex x Stress Group interaction, *F*_6,207_ = 2.426; *p* = 0.027; Figure 4D, E). To quantify the change in saccharine consumption across the 4 conditioning sessions, a percent decrease was calculated between session 1 (i.e., pre-conditioning baseline) and the average consumption in sessions 2-4 (Figure 1F). We found that males across all stress groups eventually acquired alcohol CTA after 4 conditioning sessions (main effect of Treatment *F_1,52_ =* 32.990, *p* < 0.001). Females generally also acquired aversion to alcohol (main effect of Treatment *F_1,46_* = 9.10, *p* = 0.004), but this effect was driven by female Non-Avoiders (*F_1,18_* = 25.106, *p* < 0.001).

**Figure 2:**
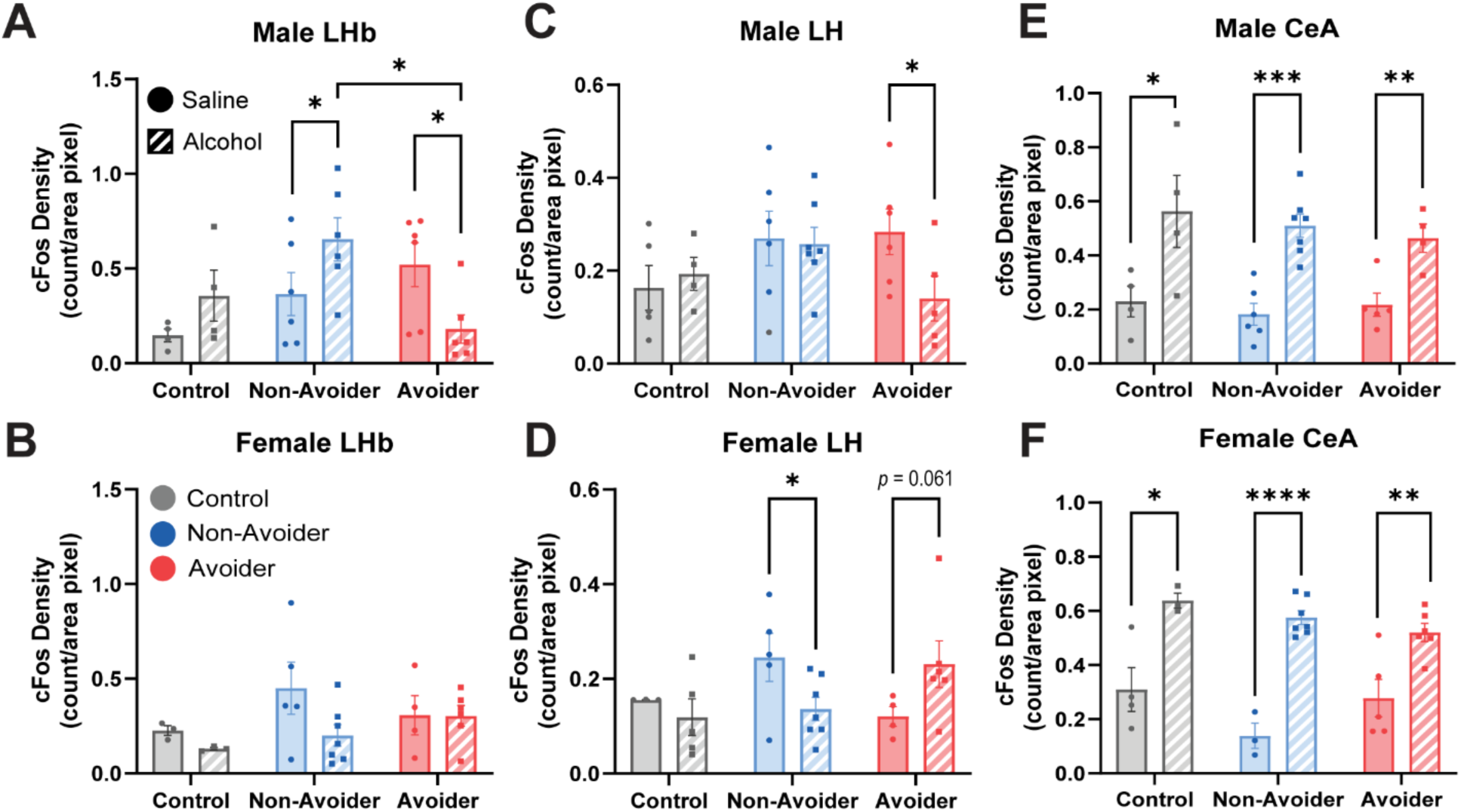
Alcohol blunts activity in the LHb of Male Avoiders and increases activity in the CeA after stress and 1.5 g/kg ethanol i.p. injection. **A, C, E.** cFos density (count of positive cFos expressing cells per area measured in pixels) of males separated by stress group (Control, Non-Avoider, and Avoider) and treatment with each stress group (Saline or Alcohol). **B, D, F.** cFos density (count of positive cFos expressing cells per area measured in pixels) of females separated by stress group (Control, Non-Avoider, and Avoider) and treatment with each stress group (Saline or Alcohol).

**Figure 3:**
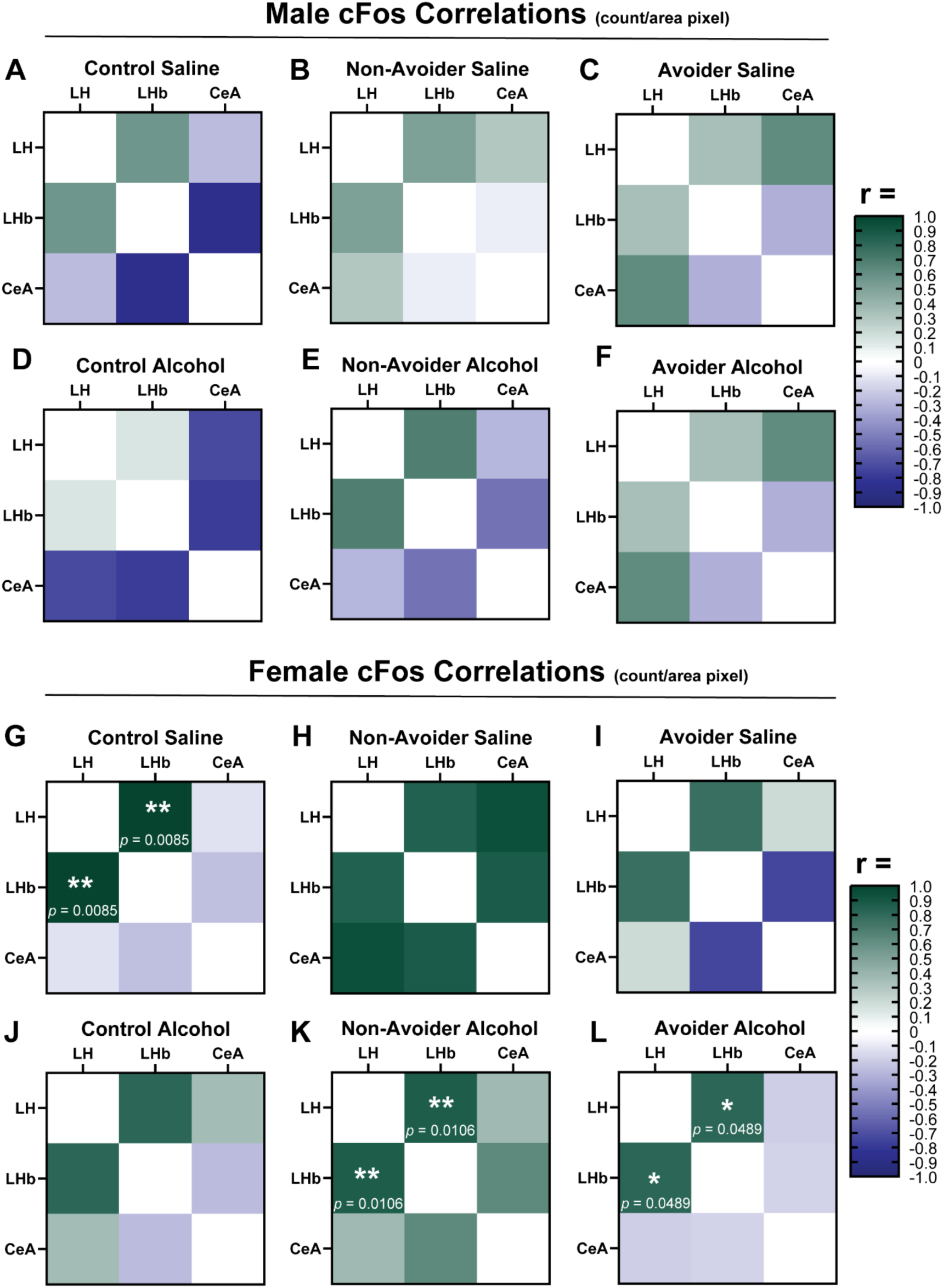
Correlations of cFos density between key regions. Heat maps of cFos density (measured by count per area in pixels) of the LH, LHb, and CeA divided by sex, stress group, and treatment of alcohol or saline. In each sex, the cFos densities between brain areas were compared within stress group and treatment. Correlations between cFos density in the LH and LHb were seen in the Control Saline, Non-Avoider Alcohol, and Avoider Alcohol groups.

**Figure 4:**
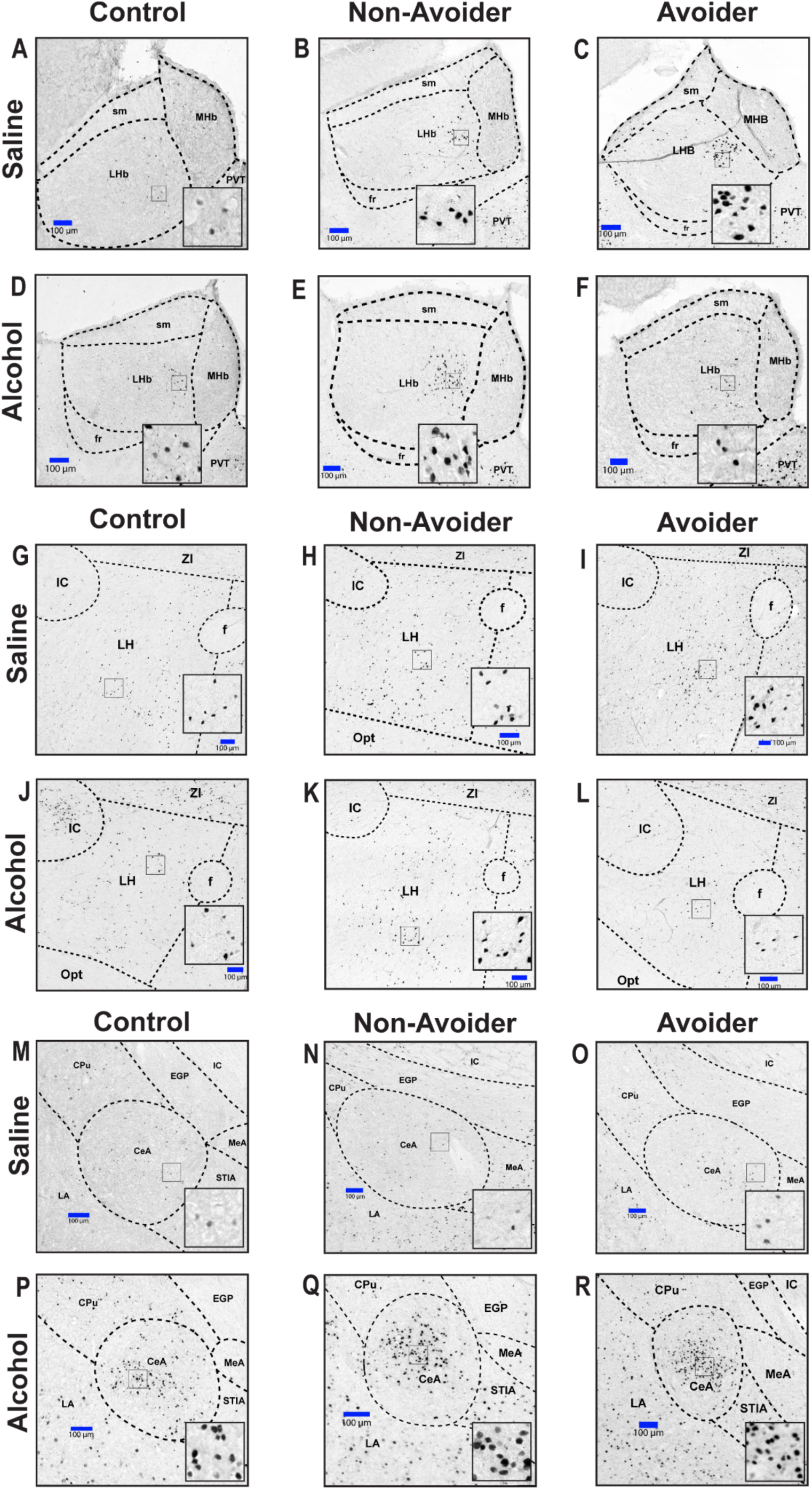
Representative images of regions quantified for cFos. Sample images of the LHb, LH, and CeA from male rats reflecting all experimental conditions: Control, Non-Avoider, or Avoider treated with Saline or Alcohol. **Abbreviations:** MHb: medial habenula, sm: stria medullaris of the thalamus, PVT: paraventricular thalamic nucleus, fr: fasciculus retroflexus, IC: internal capsule, f: fornix, ZI: zona incerta, Opt: optic tract, EGP: external globus pallidus, MeA: medial amygdala, LA: lateral amygdala, STIA: striatum, CPu: caudate putamen (striatum).

To measure the relationship between avoidance of stress-paired contexts and magnitude of alcohol CTA across all 4 sessions, we performed correlational analyses between avoidance scores and percent decrease in saccharin consumption in male and female stress-exposed rats. While there was no significant relationship between these measures in saline or alcohol-injected males (Figure 1G), greater avoidance tended to predict less alcohol CTA in alcohol-injected female rats (R^2^ = 0.1630, r = 0.4037, *p* = 0.0865; Figure 1H), supporting the idea that female Avoiders are less sensitive to alcohol’s aversive effects even after 4 conditioning sessions.

### 3.2 Sex-and stress group-specific differences in cFos expression following alcohol challenge

The goal of this study was to investigate the effects of an aversive dose of alcohol on cFos expression in the LHb, LH, and CeA of unstressed animals and stressed Avoiders vs. Non-Avoiders. Rats were subjected to the 4-day predator odor place conditioning procedure and indexed for avoidance. On the 5^th^ day (i.e., 48h after predator odor or no odor exposure), animals were injected with either 1.5g/kg of alcohol or saline 90 minutes prior to euthanasia. Their brains were fixed and stained for cFos via IHC. The results of cFos quantification and analysis are as follows.

#### Alcohol increases cFos expression in the LHb of male Non-Avoiders but lowers cFos expression in male Avoiders

In male rats, challenge with an aversive, CTA-producing dose of alcohol (1.5 g/kg) increased cFos density in the LHb of Non-Avoiders but decreased cFos density in Avoiders, compared to saline injection (Figure 2A; 3-way sex x stress group x treatment interaction, *F_2,59_* = 5.799, *p* = 0.006; treatment x stress group interaction within males, *F_2,31_* = 5.584, p=0.010; Alcohol vs. Saline effects in Non-Avoiders [p=0.051] and Avoiders [*p* = 0.017]). A direct comparison of LHb cFos density between male Avoiders, Non-Avoiders, and Controls also showed that alcohol significantly diminished cFos expression in Avoiders, compared to Non-Avoiders (Tukey’s post-hoc, *p* = 0.013). In females (Figure 2B), there was not a significant interaction between treatment and stress group (*F_2,27_* = 1.220, p=0.314), or main effect of stress group (*F* = 1.370, *p* = 0.124) or treatment (*F* = 2.562, *p* = 0.124).

#### Alcohol lowers cFos expression in the LH of male Avoiders and female Non-Avoiders

In male rats, a challenge with an aversive dose of alcohol (1.5 g/kg) diminished cFos expression in the LH of Avoider rats (Figure 2C. 2D; 3-way sex x stress group x treatment interaction, *F_2,62_ =* 4.769, *p* = 0.013; Alcohol vs. Saline effect in male Avoiders [*p* = 0.035]). In females, alcohol challenge decreased cFos expression in Non-Avoiders (2-way stress x treatment interaction, *F_2,29_ =* 4.198, *p =* 0.027; Alcohol vs Saline effect in female Non-Avoiders [*p* = 0.029]). Additionally, alcohol challenge mildly increased cFos expression in female Avoiders (*p* = 0.061).

#### Alcohol increases cFos expression in the CeA across sexes and stress groups

An aversive dose of alcohol increases cFos expression in the CeA of all experimental animals. Males and females were analyzed separately to reflect the analysis of the previous brain regions mentioned despite the lack of a significant interaction between sex, stress group, and treatment (*F_2,57_* = 0.472, *p* = 0.627). There were no significant interactions between stress group and treatment in either males (Figure 2E; *F_2,29_* = 0.293, *p* = 0.749) or females (Figure 2F; *F_2,27_* = 2.502, *p* = 0.105). In males, there was a significant effect of treatment in Non-Avoiders (*F_1,12_* = 29.996, *p* = 1.929x10^-4) and Avoiders (*F_1,8_* = 13.444, *p* = 0.008). A one-sided t-test showed an effect of treatment in male Controls (p=0.050). In males, alcohol greatly increased cFos expression in the CeA across all groups (Alcohol vs Saline effect across stress conditions; *F_1,29_* = 34.901, *p* = 4.277x10^-6; Alcohol vs Saline effect in Controls *p =* 0.050, Non-Avoiders *p* = 1.9x10^-4, and Avoiders *p* = 0.008), and the same was observed in females (Alcohol vs Saline effect across stress conditions, *F_1,27_* = 67.079, *p* = 3.988x10^-8^; Alcohol vs Saline effect in Controls, *p* = 0.020, Non-Avoiders, *p =* 9.308x10^-5^, and Avoiders, *p* = 0.009).

#### Alcohol increases correlations between LH and LHb cFos densities in stressed female rats

We performed correlational analyses of cFos cell counts in the LHb, LH, and CeA to explore potential relationships that were either recruited or disrupted by an acute challenge with an aversive dose of alcohol. Correlational analyses of cFos densities in male rats showed that none of these cell counts were significantly correlated (Figure 3A-F). However, in Non-Avoiders (Figure 3B, E), alcohol increased the cFos correlation between the LH and LHb from an r value of 0.504 to 0.686. In female rats, while alcohol weakened the correlation between LH and LHb cFos expression in unstressed Controls (Figure 3G, Saline: r = 0.991, *p* = 0.0085; Figure 3J, Alcohol: r = 0.814, *p* = 0.186), alcohol strengthened LH-LHb cFos correlations in stress-exposed Non-Avoiders (Figure 3H, Saline: r =0.85, p=0.0684; Figure 3K, Alcohol; r =0.872, *p* = 0.0106) and Avoiders (Figure 3I, Saline: r = 0.769, *p* = 0.231; Figure 3L, Alcohol: r = 0.814, p=0.0489).

### 3.3 Blood alcohol concentrations reflect injection dose

The purpose of this experiment was to assess blood alcohol concentrations (BACs) produced by intraperitoneal injection of 1.5 g/kg alcohol, a dose that produced CTA and changes in cFos expression in the present studies. Male and female rats were injected with either 0.5, 1.5, or 2.0 g/kg alcohol and blood samples were collected at 30- and 90-min post-injection. 1.5 g/kg alcohol produced BACs of 142.425 ± 11.875mg/dl in males (Figure 5A) and 168.7 ± 6.400 mg/dl in females 30 minutes after injection (Figure 5B). BACs were 135.925 ± 0.925 and 167.9 mg/dl for females and males, respectively, 90 minutes after injection. Figure 5 shows all BAC measurements taken from male and female rats injected with all three alcohol doses.

**Figure 5:**
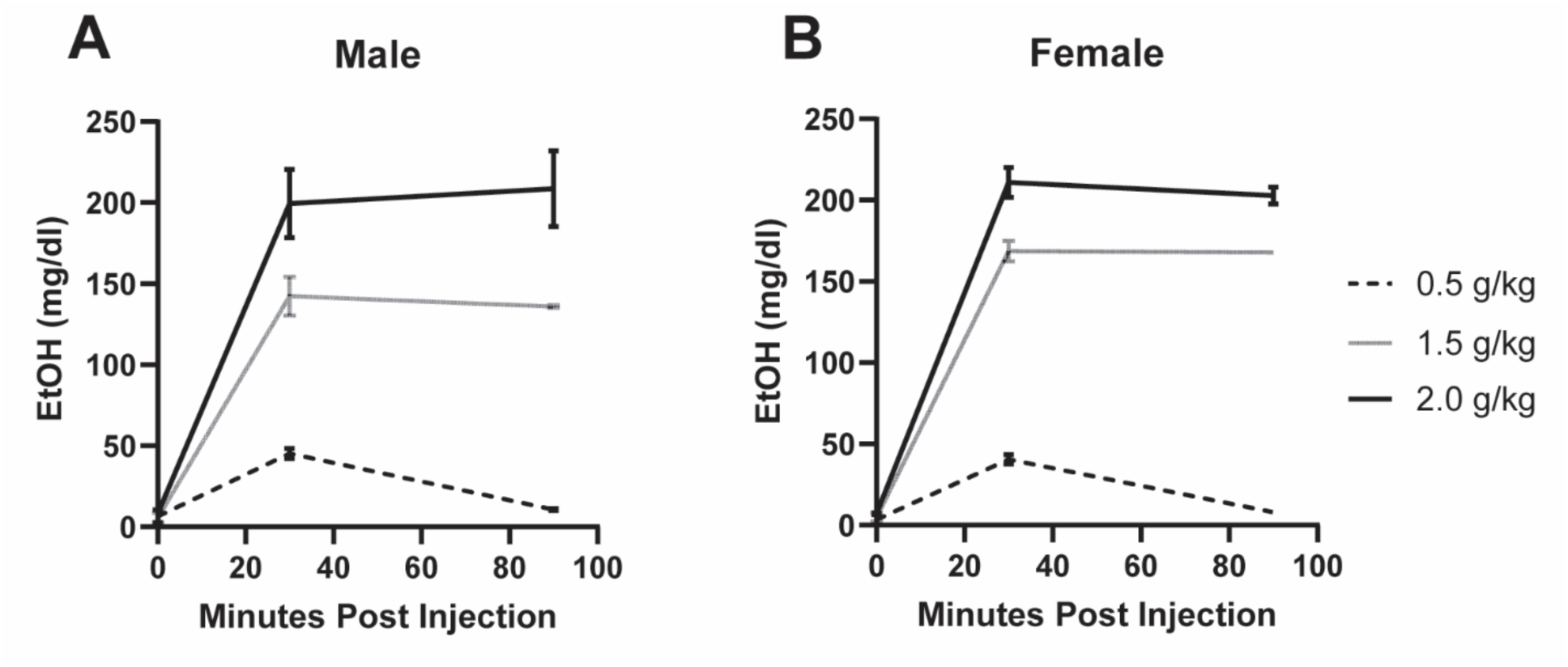
Blood Alcohol Concentrations. **A**. The concentration of ethanol (mg/dl) in serum of males before injection and 30 and 90 minutes after injection. **B.** The concentration of ethanol (mg/dl) in serum of females before injection and at 30 and 90 minutes after injection.

### 3.4 Avoider females exhibit hyperactive behavior during predator odor conditioning Posttest

The goal of this analysis was to more comprehensively quantify anxiety-like behaviors in Avoider and Non-Avoider rats during expression of avoidance or non-avoidance of stress-paired contexts. Rats exposed to predator odor stress are typically stratified into Avoider and Non-Avoider behavioral phenotypes based on the change in time spent in the odor-paired context between the Pretest and the Posttest (Edwards et al., 2013; Albrechet-Souza and Gilpin, 2019). In general, male and female Avoiders spend less time in the odor-paired context than Non-Avoiders, as demonstrated by heat maps from representative individuals (colors represent proportion of total duration spent in each location, red indicates greater amount of time, Figure 6A). Here, 50% of male rats (n = 20) and 44% of female rats (n = 16) were classified as Avoiders following predator odor place conditioning (Figure 6B).

**Figure 6.**
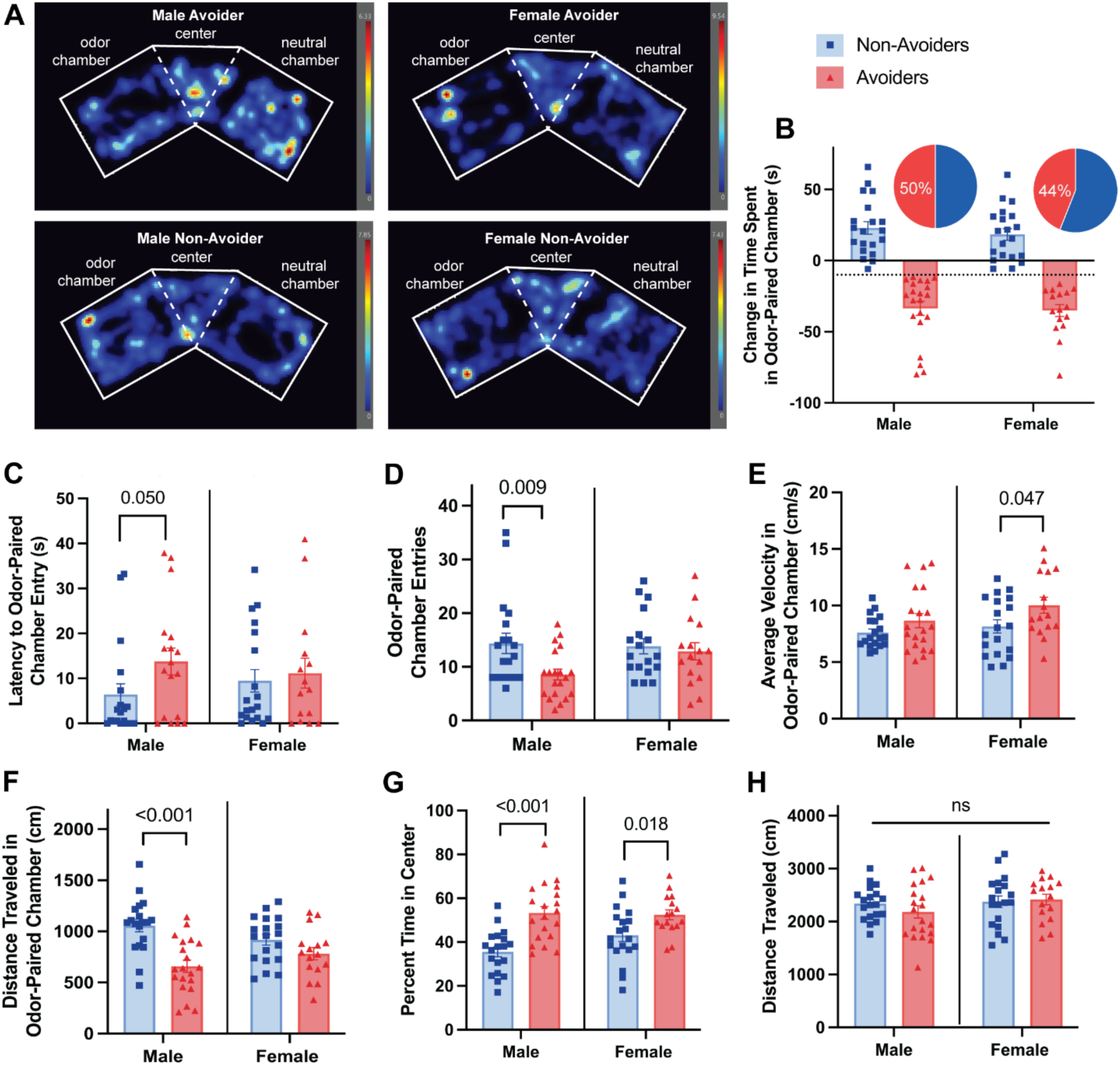
Hyperactive exploration behavior in Avoider females during predator odor CPA. **A**. Representative heat maps for one rat in each stress group and sex. Color scale represents the proportion of total duration (5 minutes) in designated location, red is greater time. **B.** Stress group outcomes based on 4-day predator odor CPA for males and females (male Avoiders = 20; female Avoiders = 16). **C.** Latency to enter the odor-associated context during Posttest for all stressed rats. **D.** Number of entries into odor-paired context for stressed rats. **E.** Average velocity within the odor-paired context. **F.** Total distance traveled within the odor-paired chamber. **G.** Percentage of time outside of odor-paired context that was spent in the center zone (proximal to odor-paired context). **H.** Total distance traveled within the entire apparatus during avoidance Posttest.

Within stressed rats, there were marked differences in the patterns of approach/avoidance and exploration behavior relative to the odor-paired context. Males and females were analyzed separately to illuminate stress group differences within each sex. Analysis of latency to first enter the odor-paired context was not normally distributed for males or females. Mann-Whitney U Tests revealed that Avoider males were more latent to enter the odor-paired context compared to Non-Avoiders (U = 113.0, n = 38, *p* = 0.050) while there were no stress group effects on females (Figure 6C). Similarly, while Avoider males had significantly fewer number of entries into the odor-paired context than Non-Avoider males (F_1,37_ = 7.631, *p*=0.009), females in both stress groups entered nearly the same number of times (Figure 6D). Within the odor-paired context, males had similar average velocity patterns, regardless of stress group, while female Avoiders displayed greater average velocities compared to Non-Avoiders indicating hyperactive exploration (F_1,33_ = 4.270, *p*=0.047; Figure 6E). Importantly, all females moved similar distances while in the odor-paired chamber, while Avoider males moved less distance than Non-Avoiders, suggesting that even though Avoider females spent less time in the odor-paired chamber, their locomotion was indeed accelerated and hyperactively exploratory (F_1,37_ = 21.330, *p*<0.001; Figure 6F). When stressed rats were not in the odor-paired context, Avoiders of both sexes spent a larger percentage of time in the center zone compared to Non-Avoiders, essentially spending more time proximal to the odor chamber (males F_1,37_=22.348; *p*<0.001; females F_1,33_=6.178; *p*=0.018; Figure 6G). Finally, total distance traveled in the entire arena during the Posttest was consistent across stress groups for both males and females, indicating that behavioral differences were not driven by baseline locomotion (Figure 6H). In all, this provides greater insight into the stress-group differences of other anxiety-like behaviors and avoidance coping behaviors during the expression of CPA. Of particular note is the pattern of female Avoiders showing a more active exploration compared to males, but ultimately comparable avoidance durations.

## 4. Discussion

The described experiments support the hypothesis that a diminished sensitivity to alcohol’s aversive effects is a key contributor to escalated alcohol consumption in Avoider rats following stress exposure. We showed that while a single alcohol challenge at a dose of 1.5 g/kg (producing BACs of ∼150 mg/dl) was sufficient to produce conditioned aversion to a novel tastant (i.e., CTA to saccharin solution) in Non-Avoider rats, Avoiders were resistant to this phenomenon. Although male Avoider rats did eventually develop alcohol-induced CTA after repeated conditioning sessions, female Avoiders seem to have remained resistant to alcohol’s aversive effects. In other words, Non-Avoider rats across both sexes appear to have protective mechanisms that promote sensitivity to alcohol’s aversive effects. An aversive dose of alcohol produced different patterns of cFos activity in the lateral habenula and lateral hypothalamus of Avoiders vs. Non-Avoiders. While alcohol induced cFos expression in LHb cells of male Non-Avoiders, LHb cFos expression was blunted by alcohol in male Avoiders compared to their saline-injected counterparts. The cFos activity in the LH was similarly blunted by alcohol challenge in male Avoider rats. Interestingly, the pattern of cFos expression in LH was flipped in females, such that Non-Avoiders treated with alcohol have blunted cFos expression compared to saline treated Non-Avoiders.

In humans, individual differences in sensitivity to the interoceptive effects of alcohol play a role in determining risk for developing AUD. In general, lower sensitivity to the negative effects of alcohol predicts higher risk for developing AUD (Schuckit 2004, 2014; King 2011, 2014; Koob 2013a, 2013b) In animal models, there is a similar inverse relationship between sensitivity to alcohol’s acute aversive effects and drinking propensity. For example, rodents with lower sensitivity to alcohol-induced CTA have higher voluntary ethanol intake (Broadbent et al 2002; Green and Grahame 2008). Similarly, mice bred for high alcohol preference are resistant to developing alcohol induced CTA compared to low alcohol preferring mice strains (Chester et al., 2003). Further analysis of genetic variations in sensitivity of mice to alcohol’s aversive effects shows that reduced sensitivity is linked to a predisposition for heightened alcohol drinking behavior (Cunningham 2013, 2019). In rats, a single inescapable exposure to predator odor (potentially ‘traumatic’) stress increases alcohol self-administration and alcohol drinking in animals classified as Avoiders but not in Non-Avoiders (Edwards et al., 2013; Weera et al., 2023; Bonauto et al., 2025). In the study at hand, stress-exposed Non-Avoiders developed alcohol-induced CTA after a single conditioning session while Avoiders did not, indicating that Non-Avoider rats have increased sensitivity to the aversive effects of alcohol. Given that Non-Avoiders do not escalate their alcohol drinking after stress, their heightened sensitivity to alcohol’s aversive effects may serve as a moderator of their alcohol intake. All rats across all groups eventually developed alcohol-induced CTA after multiple (four) conditioning sessions, but alcohol CTA in females remained driven by Non-Avoiders.

The lateral habenula plays a key role in aversion-related behavior, learning (Matsumoto and Hikosaka 2007, 2009), and addiction (Velasquez et al., 2014). Neurons in the LHb are excited by aversive stimuli, such as alcohol and foot shock stress, and inhibited by rewarding stimuli (Matsumoto and Hikosaka 2007, 2009; Wang et al 2017; Stamatakis & Stuber 2012). Lesions of the LHb reduce alcohol-induced CTA and prevent yohimbine—a stressor—from reinstating alcohol-seeking behavior (Haack et al., 2014). Similarly, expression of alcohol-induced CTA increases neuronal activity in the LHb, implicating this nucleus in signaling alcohol’s aversive effects (Glover et al., 2016). Downstream of the habenula, the ability of LHb neurons to signal aversion is mediated by their projections to the rostromedial tegmental nucleus (Stamatakis and Stuber 2012). In the present study, an alcohol challenge increased cFos expression in stress-exposed male Non-Avoider rats (and, to a lesser degree, unstressed controls), in support of existing literature showing that LHb neurons are activated in response to the aversive effects of alcohol (Tandon et al., 2017; Haack et al., 2014). In contrast, Avoiders treated with alcohol had significantly blunted cFos expression in the LHb. These dichotomous effects of alcohol on LHb activity in male Avoiders vs. Non-Avoiders are likely to contribute to differences in sensitivity to alcohol’s aversive effects, and perhaps to general differences in aversion signaling in these animals. In females, alcohol surprisingly did not alter cFos expression in the LHb. Although most of the studies mentioned regarding the role and functioning of the LHb are from male subjects, there is some work in females that show that the LHb is indeed important for aversion signaling in these animals. For instance, restraint stress has been shown to increase LHb activity in female mice (Kim and Chung 2021). With regards to alcohol responsivity, one possibility is that LHb cells may have a left-shifted alcohol dose-response curve, such that these cells may be activated by lower doses of alcohol and inhibited by higher doses. Future work will test a range of alcohol doses on LHb cell activity in female Avoiders and Non-Avoiders.

The lateral hypothalamus (LH) is a brain area that provides dense glutamatergic inputs to the LHb (Poller et al., 2013; Groos et al., 2025). The LH is well known to mediate physiological and behavioral stress responses and serves as a key modulator of consummatory and approach vs. avoidance behaviors (Bonnavion et al 2016). A major pathway by which LH neurons signal aversion and produce avoidance behavior appears to be via the LHb. Indeed, stimulation of LH projections to the LHb has been shown to support avoidance behaviors (Lecca et al., 2017; Lazaridis et al., 2019). However, it is unclear if this circuit, and if the LH in general, are involved in signaling alcohol’s aversive effects. In this study, an aversive dose of alcohol diminished cFos expression specifically in the LH of male Avoider and female Non-Avoider rats. Given the complexity and heterogeneity of LH circuitry, general cFos quantification lacks specificity to fully disentangle sex-, stress-, and alcohol-related effect in this experiment. However, examination of the correlation between cFos activity in the LH and LHb in this study reveals more about the effects of alcohol on LH-LHb circuitry. Correlational analyses of LH and LHb cFos expression suggest that an aversive dose of alcohol strengthened the functional connectivity between the LH and LHb in male Non-Avoiders, but not in Avoiders. Additionally, LHb and LH neural activity was positively correlated in alcohol-treated Non-Avoider and Avoider females, but not in Controls. Thus, alcohol could blunt the functional connectivity between the LH and LHb in controls, while enhancing it in stressed female animals. The strength of the LH-LHb functional connectivity in Non-Avoiders could explain their higher CTA sensitivity compared to the other females. The LH-LHb circuit seems to vary at the individual level and could be an important target for preventing alcohol misuse.

Indeed, LH projections to LHb are necessary for regulating alcohol drinking. (Sheth et al., 2017). Stimulation of LH-LHb circuit has been shown to signal aversion while inhibition of the circuit is rewarding (Stamatakis et al., 2016). Therefore, strong relationship between these regions is expected as observed in females after alcohol treatment. Another key mechanism within the LH is the orexin system, as orexin is uniquely produced in this region and the system plays a critical role in stress, reward, and addiction pathways (Giardino & de Lecea 2014; Boutrel & de Lecea 2008; Boutrel et al., 2005; Aston-Jones et al., 2010; Hopf et al., 2020). The strength of orexin neuron activation has been found to correlate with alcohol seeking (Moorman et al., 2016), while stimulation of orexin positive LH cells produces real time place preference (Giardino et al 2018). Importantly, orexin 1 receptor antagonism has been shown to reduce alcohol consumption, seeking, self-administration, and relapse (Moorman et al., 2017; Anderson et al 2014) and prevent stress-induced reinstatement of alcohol seeking (Richards et al., 2008), implicating this system in LH-mediated alcohol behaviors. We are currently using circuit-based techniques to test the role of LHb-projecting LH neurons and the orexin system in alcohol aversion.

The central amygdala (CeA) is part of the extended amygdala and is responsible for integrating sensory information related to alcohol and stress, instructing behavioral and physiological responses (Gilpin et al., 2015; Ventura-Silva et al., 2013). The CeA is recruited in response to acute alcohol treatment and by predator odor stress in male Avoider rats (Weiner and Valenzuela, 2006; Zhou et al., 2006; Roberto et al., 2003; Weera et al., 2020,). In the current study, alcohol activated the CeA in all stress groups and sexes compared to animals treated with saline. These results suggest that alcohol recruits the CeA regardless of prior stress experience; however, future studies should consider possible stress-induced changes in activity of different cell types within the CeA. A relevant system to consider is CRF, as CRF1-expressing cells in the CeA projecting to the LH are more active in Avoiders. Additionally, we have identified sex and stress group differences in CRH-system gene expression in the LH which the CeA could be influencing (Bonauto et al., 2025), underscoring the need for further investigation of the connectivity of these complex regions. Although no stress or sex effects in the CeA were observed, the CeA may still influence LH function in alcohol aversion.

To better understand the differences between Avoider and Non-Avoider behavioral phenotypes, we measured additional behaviors during post-stress avoidance of the odor context. This analysis revealed a pattern of hyperactive, escape-like behavior in female Avoiders that was not present in female Non-Avoiders or any of the male stressed rats. Male Avoiders were slower to enter the odor-paired context, made fewer entries into the context, and had movement velocities in the odor chamber comparable to Non-Avoiders, aligning with expected patterns of avoidance.

In contrast, female Avoiders showed no difference in latency or number of entries compared to Non-Avoiders yet had greater movement velocities in the odor-paired chamber, indicating rapid, hypervigilant exploration of the context despite ultimate avoidance. Avoider females also traveled nearly the same distance in the odor context as Non-Avoiders, despite spending less time there. All Avoiders clearly remembered the odor-paired context, because both male and female Avoiders spent more time in the center area proximal to the odor-paired chamber (as a percentage of time not in the odor context). Thus, all Avoiders somewhat investigated the context without crossing the threshold. Additionally, these movement patterns were not a function of locomotion differences, as the distance traveled within the entire apparatus was similar across all groups.

Hyperactive, escape-like behaviors have been previously observed in stressed female rats. In traditional assays of anxiety-like behavior such as the elevated plus maze or open field, female rats travel further distances than males and enter anxiogenic areas more frequently, with no effect of estrous cycle phase (Knight et al., 2021; Scholl et al., 2019). In fear conditioning studies, some female rats respond with darting at high velocities, while males are less likely to do so (Gruene et al., 2015) suggesting that patterns of conditioned responses to stressors can differ by sex. Using predator odor stress place conditioning, few studies have considered sex differences in addition to avoidance outcomes. Some studies have noted that a greater number of females become Avoiders compared to males (Albrechet-Souza and Gilpin, 2019; Bonauto et al., 2025), however, most studies have only utilized males. Additionally, we previously observed that predator odor stress induced hypervigilant, anxiety-like behavior in Avoider females up to 1 month after stress, measured by aborted attempts to enter the elevated-plus maze open arm, (Bonauto et al., 2025). However, another group found that all predator odor stressed females exhibited blunted acoustic startle response compared to unstressed females, this suggests hyperarousal after stress may be time dependent and test-specific (Albrechet-Souza et al., 2020). In all, these studies suggest that predator odor stress induces sex-specific differences in post stress behavior. Here we provide evidence that these differences emerge just 24 hours after stress, during expression of avoidance behavior in predator odor place conditioning Posttest session. Given the established role of the LH and LHb in avoidance and anxiety-like behavior (Matsumoto and Hikosaka 2007, 2009; Bonnavion et al., 2016), stress-induced dysregulation of these regions and recruitment during avoidance may drive post stress behaviors and contribute to sex differences. Future work should consider sex and stress group differences in both behavior and neural recruitment during predator odor stress exposure itself.

Overall, we identified key stress-induced changes in sensitivity to alcohol-induced CTA following predator odor stress in males and females. We demonstrate that alcohol-induced neural recruitment differs based on avoidance response to stress in the LHb, LH, and CeA with only a single dose of alcohol. Alcohol blunts LHb activity of male Avoiders but elevates it in male Non-Avoiders. In females, LH and LHb activity are strongly correlated in a stress- and alcohol-dependent manner. Finally, we show that males and females have distinct behavioral patterns during avoidance indexing, indicating a hyperactive stress response that is unique to female Avoiders. Future research should further explore the LH-LHb circuit, given strong evidence of its involvement in alcohol aversion after a single dose. More research is needed to investigate sex differences, particularly in females within the LH and LHb.

